# Reticulate Evolutionary History in a Recent Radiation of Montane Grasshoppers Revealed by Genomic Data

**DOI:** 10.1101/2021.01.12.426362

**Authors:** Vanina Tonzo, AdriÀ Bellvert, Joaquín Ortego

**Author notes:** Author for correspondence: Vanina Tonzo, Estación Biológica de Doñana, EBD-CSIC, Avda. Américo Vespucio 26, E-41092 Seville, Spain.

## Abstract

Inferring the ecological and evolutionary processes underlying lineage and phenotypic diversification is of paramount importance to shed light on the origin of contemporary patterns of biological diversity. However, reconstructing phylogenetic relationships in recent evolutionary radiations represents a major challenge due to the frequent co-occurrence of incomplete lineage sorting and introgression. In this study, we combined high throughput sequence data (ddRADseq), geometric morphometric information, and novel phylogenetic inference methods that explicitly account for gene flow to infer the evolutionary relationships and the timing and mode of diversification in a complex of Ibero-Maghrebian montane grasshoppers of the subgenus *Dreuxius* (genus *Omocestus*). Our analyses supported the phenotypic distinctiveness of most sister taxa, two events of historical introgression involving lineages at different stages of the diversification continuum, and the recent Pleistocene origin (< 1 Ma) of the complex. Phylogenetic analyses did not recover the reciprocal monophyly of taxa from Iberia and northwestern Africa, supporting overseas migration between the two continents during the Pleistocene. Collectively, these results indicate that periods of isolation and secondary contact linked to Pleistocene glacial cycles likely contributed to both allopatric speciation and post divergence gene flow in the complex. This study exemplifies how the integration of multiple lines of evidence can help to reconstruct complex histories of reticulated evolution and highlights the important role of Quaternary climatic oscillations as a diversification engine in the Ibero-Maghrebian biodiversity hotspot.

## Introduction

Recent evolutionary radiations have traditionally received much attention because the signatures of speciation events have not been fully erased by time and, thus, provide the potential to infer processes from fine-scale patterns of genetic and phenotypic variation (Shaw and Danley 2003; Shaffer and Thomson 2007; Knowles and Chan 2008). Phylogenies provide essential tools to infer the processes responsible for speciation, investigate trait evolution, and discern among alternative biogeographic scenarios (Barraclough et al. 1998; Knowles and Chan 2008). Inferring the mode and timing of speciation is crucial to reconstruct the diversification process and unravel the origin of contemporary patterns of biological diversity. However, reconstructing phylogenetic relationships among recently diverged species can be extremely challenging. One of the main issues is the frequent co-occurrence of incomplete lineage sorting and introgression (Maddison 1997; Nichols 2001; Edwards 2009).

Although phylogenetic relationships among species have been typically represented as bifurcating branches (Haeckel 1866; Felsenstein 2004), which implicitly assumes that diversification occurred without reticulation (Coyne and Orr 2004; Mallet 2007), there are multiples examples of gene flow among independently evolving taxa (Feder et al. 2012; Harrison and Larson 2014; Burbrink and Gehara 2018; Blair et al. 2019). Thus, failing to account for post-divergence gene flow when estimating evolutionary processes may produce statistical inconsistencies, incorrect phylogenies, inaccurate estimates of key demographic parameters, and wrong biogeographic inferences (Solís-Lemus et al. 2017; Burbrink and Gehara 2018; Flouri et al. 2018).

Speciation events driven by high amplitude climatic variations in the Middle and Late Pleistocene (774 ka to 10 ka), are among the best-known examples of recent diversification processes (Roy et al. 1996; Flantua and Hooghiemstra 2018). Repeated range expansions and contractions driven by Quaternary glacial cycles have extraordinarily contributed to the diversification of montane and alpine biotas (Hewitt 1996; Shepard and Burbrink 2008; Sandel et al. 2011; Wallis et al. 2016). Interglacial periods pushed cold-adapted lineages from mid and low latitude regions to shift their distributions towards high elevations to satisfy their specific habitat and climate niche requirements, leading to range fragmentation and divergence in interglacial refugia (e.g., DeChaine and Martin 2005; Djamali et al. 2012). Conversely, glacial periods forced downslope migrations in montane organisms, which likely experienced net range expansions, colonization of new suitable habitats in lowlands and secondary contact and admixture among closely related lineages (Hewitt 1990; Excoffier et al. 2009; Marko and Hart 2011). Glacial advances also contributed to allopatric divergence in alpine biotas, particularly those inhabiting extensively glaciated and topographically complex regions where distributional ranges got severely fragmented by ice caps and valley glaciers and populations likely became confined to highly isolated ice-free refugia (Wallis et al. 2016). Isolation periods contributed to genetic and phenotypic differentiation, fueling allopatric adaptive (i.e., divergent natural selection) and non-adaptive (i.e., genetic-drift) lineage divergence and/or reinforcing existing species boundaries (Hewitt 1996, 1999; Czekanski-Moir and Rundell 2019). If reproductive isolation did not evolve while in refugia, secondary contact during range shifts resulted in the collapse of formerly distinct lineages (i.e., speciation reversal; Kearns et al. 2018; Maier et al. 2019), introgressive hybridization (e.g., Salzburger et al. 2002; Schweizer et al. 2019), or even contributed to complete the speciation process via reinforcement of reproductive isolation (Butlin and Hewitt 1985; Hewitt 1996; Nevado et al. 2018). For these reasons, Pleistocene glacial cycles have been considered to both promote range fragmentation and allopatric speciation (Knowles 2000) and inhibit speciation through genetic homogenization (Zink and Slowinski 1995; Klicka and Zink 1997).

The Iberian Peninsula and western Maghreb regions present a rich biodiversity and an alike species composition due to their close geographical proximity, similar climatic and ecological conditions, complex topography, and a geological history that has led to multiple episodes of connectivity and isolation for terrestrial biotas distributed in the two continents (Blondel and Aronson 2002; Krijgsman 2002; Meulenkamp and Sissingh 2003). As a result, this region is an important center of diversification for numerous organism groups and considered a hotspot for animal and plant biodiversity (Rodríguez-Sánchez et al. 2008; Myers et al. 2020). The re-opening of the Strait of Gibraltar at the beginning of the Pliocene led to the loss of the last intercontinental land connection stablished during the desiccation of the Mediterranean Basin in the Messinian Salinity Crisis (Krijgsman 2002; Husemann et al. 2014), a phenomenon representing the starting point for the diversification of many lineages whose distributional ranges resulted fragmented under the new geographic setting (e.g., Veith et al. 2003; Faille et al. 2014). However, empirical evidence has also supported that the shortening of coastline distances during Pleistocene glacial periods facilitated fauna exchanges and gene flow between southern Europe and North Africa (Agustí et al. 2006; Carranza et al. 2006; Graciá et al. 2013). In this context, resolving the phylogenetic relationships among Ibero-Maghrebian species complexes and estimating their timing of divergence is essential to unravel whether their origin is linked to Pleistocene range expansions/contractions (e.g., Knowles 2000) and sea-level low stands (e.g., Graciá et al. 2013) or, rather, compatible with a protracted history of diversification dating back to the late Miocene (e.g., Hidalgo-Galiana and Ribera 2011; Faille et al. 2014).

Here we focus on the Ibero-Maghrebian subgenus *Dreuxius* Defaut, 1988 (genus *Omocestus* Bolívar, 1878), a complex of montane grasshoppers (Orthoptera: Acrididae) currently comprised by eight species distributed in the Iberian Peninsula (5 species) and northwestern Africa (3 species) (Tonzo et al. 2019; Cigliano et al. 2020). Most taxa present allopatric distributions and form isolated populations at high elevations in different mountain systems (Tonzo et al. 2019, 2020; Cigliano et al. 2020; Fig. 1). The only exceptions are the Iberian *O. minutissimus* (Brullé 1832) and the Maghrebian *O. lecerfi* Chopard 1937, which present wider elevational ranges and geographic distributions partially overlapping with the rest of Iberian and northwestern African species of the complex, respectively, and with which they often form sympatric populations (Clemente et al. 1990; Cigliano et al. 2020; Tonzo et al. 2020). All taxa within the complex are predominantly graminivorous and their distributions are tightly linked to open habitats of cushion and thorny shrub formations (e.g., *Erinacea* sp., *Festuca* sp., *Juniperus* sp., *Thymus* sp.) that they use as refuge (Gangwere and Morales Agacino 1970; Clemente et al. 1990). Species within this subgenus, particularly females, are markedly brachypterous, which is expected to extraordinarily limit their dispersal capacity, reduce gene flow at short spatial scales and, ultimately, might have contributed to genetic divergence and allopatric speciation (Waters et al. 2020; e.g., Huang et al. 2020). For these reasons, this transcontinental species complex offers an ideal case study to test alternative biogeographic scenarios underlying the high rates of endemism of the region and gain insights into the proximate processes underlying species formation and patterns of phenotypic variation.

**Figure 1.**
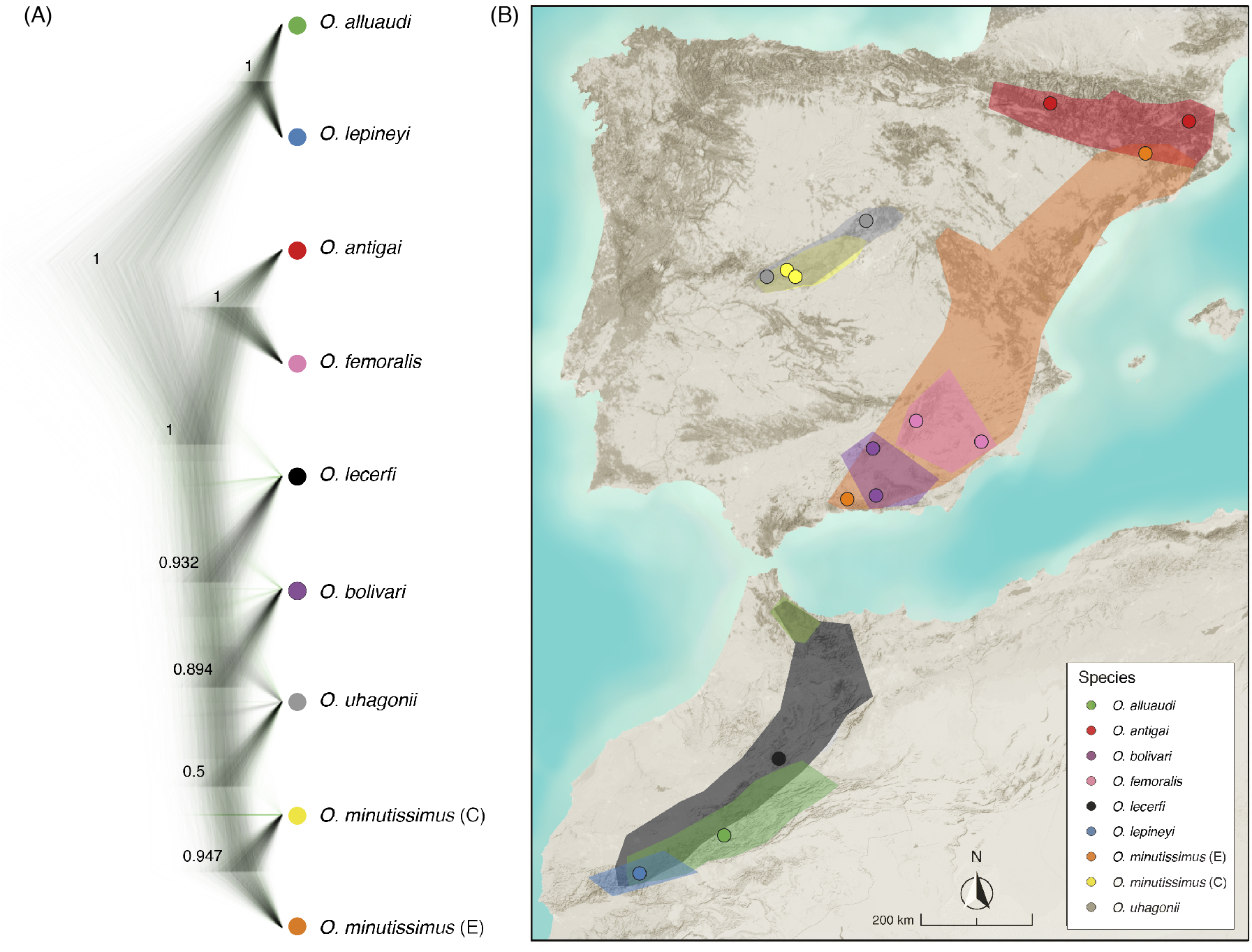
(A) Bayesian phylogenetic tree reconstructed with snapp. Posterior probabilities of clade support are indicated. (B) Approximate geographical distribution of the different species/lineages of the subgenus *Dreuxius* (shapes) and populations (dots) included in the analyses.

In this study, we integrate high throughput sequence data, geometric morphometrics, and novel phylogenetic inference methods that explicitly account for gene flow to unravel the evolutionary relationships and the timing and mode of diversification in the studied species complex. Specifically, we first generated genomic data for all species within the subgenus *Dreuxius* using a restriction-site-associated DNA sequencing approach (ddRADseq; Peterson et al. 2012) and inferred their phylogenetic relationships applying two alternative coalescent-based methods (Bryant et al. 2012; Yang 2015) and a maximum pseudolikelihood approach accounting for post-divergence gene flow (Solís-Lemus and Ané 2016). Second, we estimated species divergence times under the multispecies coalescent (MSC) model (Yang 2002; Rannala and Yang 2003) and a new implementation of the MSC model with introgression (MSCi) (Flouri et al. 2019), and evaluated the potential impact of historical gene flow on demographic parameter estimation and the inferred biogeographic history. Finally, we employed a geometric morphometric approximation (Adams and Otárola-Castillo 2013) to characterize phenotypic variation at traits of taxonomic relevance and/or putatively linked to reproductive isolation and evaluated whether such variation was shaped by a shared evolutionary history (i.e., Brownian motion under genetic drift) or departed from expectations given the phylogenetic tree, which might be indicative of selective processes acting at different stages of speciation (Gray and McKinnon 2007; Safran et al. 2013).

## Materials and methods

### Species Sampling

Between 2011 and 2017, we collected specimens representing all species of the subgenus *Dreuxius* (genus *Omocestus*) (Cigliano et al. 2020; Table 1; Fig. 1). We considered as independent lineages allopatric populations of *O. minutissimus* from central and eastern Iberia (hereafter, *O. minutissimus* C and *O. minutissimus* E, respectively), as they form distinctive genotypic and phenotypic clusters according to preliminary analyses (Cáliz 2015; Tonzo et al. 2020). Two of the taxa within the complex (*O. navasi* and *O. antigai*) have been recently synonymized on the basis of detailed genomic and phenotypic species delimitation analyses and, thus, they were considered as a single species (*O. antigai*; Tonzo et al. 2019; Cigliano et al. 2020). Whenever possible, we collected and analyzed two populations representative of the distribution range of each species/lineage (Table 1; Fig. 1). We stored specimens in 2 ml vials with 96% ethanol and preserved them at −20° C until needed for geometric morphometric and genomic analyses.

**Table 1.** Locality and geographical location (latitude, longitude and elevation) for each species and sampled population.

| Species | Locality | Latitude | Longitude | Elevation (m) |
| --- | --- | --- | --- | --- |
| *O. alluaudi* Uvarov, 1927 | Tizi n' Tirghist (Morocco) | 31.7408 | -6.3251 | 2538 |
| *O. antigai* (Bolívar, 1897) | Setcases (Spain) | 42.4276 | 2.2663 | 2145 |
| *O. antigai* (Bolívar, 1897) | Borau (Spain) | 42.6739 | -0.5793 | 1357 |
| *O. bolivari* Chopard, 1939 | Sierra de Mágina (Spain) | 37.7406 | -3.4466 | 1900 |
| *O. bolivari* Chopard, 1939 | Sierra Nevada (Spain) | 37.0964 | -3.3891 | 2446 |
| *O. femoralis* Bolívar, 1908 | Sierra de Espuña (Spain) | 37.8652 | -1.5712 | 1514 |
| *O. femoralis* Bolívar, 1908 | Poyotello (Spain) | 38.1195 | -2.6165 | 1600 |
| *O. lecerfi* Chopard, 1937 | Col du Zad (Morocco) | 32.4531 | -5.2413 | 2100 |
| *O. lepineyi* Chopard, 1937 | Jebel Oukaïmeden (Morocco) | 31.1873 | -7.8590 | 2870 |
| *O. minutissimus* (C) (Brullé, 1832) | Puerto del Pico (Spain) | 40.3458 | -5.0143 | 1340 |
| *O. minutissimus* (C) (Brullé, 1832) | Puerto de Serranillos (Spain) | 40.3067 | -4.9467 | 1600 |
| *O. minutissimus* (E) (Brullé, 1832) | Sierra de Montsec (Spain) | 42.0473 | 0.7431 | 1520 |
| *O. minutissimus* (E) (Brullé, 1832) | Sierra Tejeda (Spain) | 36.9045 | -4.0351 | 2040 |
| *O. uhagonii* (Bolívar, 1876) | Puerto de Navafría (Spain) | 40.9846 | -3.8215 | 1893 |
| *O. uhagonii* (Bolívar, 1876) | Puerto de Peña Negra (Spain) | 40.4216 | -5.3105 | 1885 |

### Genomic Library Preparation and Processing

We obtained genomic data for a total of 36 specimens representative of one or two populations per species/lineage (4 individuals per species/lineage in all cases; Table 1). Details on the preparation of ddRADseq libraries (Peterson et al. 2012) are presented in Supplementary Methods S1 available on Dryad at: https://datadryad.org/stash/share/TyAdIXuDe8IBEWGmZ7ULxR3a14OSCPGEWRgOnzwPSgA. Raw sequences were demultiplexed and pre-processed using stacks version 1.35 (Catchen et al. 2011, 2013) and assembled using pyrad version 3.0.66 (Eaton 2014). Supplementary Methods S2 available on Dryad provides all details on sequence assembling and data filtering.

### Phylogenomic Inference

We estimated species trees using two coalescent-based methods, snapp version 1.3 (Bryant et al. 2012) as implemented in beast2 version 2.4.3 (Bouckaert et al. 2014) and bpp version 4.2 (Flouri et al. 2018). SNAPP analyses are computationally highly demanding and, for this reason, we only selected two individuals per species (those with the highest number of retained reads; Supplementary Fig. S1 available on Dryad), one for each sampled population when two populations were available (i.e., 18 individuals in total). The resulting dataset retained 723 unlinked polymorphic sites shared across all taxa. We ran snapp analyses for 1,000,000 Markov chain Monte Carlo (MCMC) generations, sampling every 1,000 steps and using as gamma prior distributions for alpha and beta 2 and 2,000 values. The forward (u) and reverse (v) mutation rates were set to be calculated by snapp and we left the remaining parameters at default values. We conducted two independent runs and evaluated convergence with tracer version 1.6. We removed 10% of trees as burn-in and merged tree and log files from the different runs using LOGCOMBINER version 2.4.1. We used treeannotator version 1.8.3 to obtain maximum credibility trees and densitree version 2.2.1 (Bouckaert 2010) to visualize the posterior distribution of trees.

Complementarily, we ran bpp version 4.2 under module A01 to estimate the species tree (Yang 2015; Flouri et al. 2018). bpp program is a full-likelihood implementation of the MSC model and uses a reversible-jump Markov chain Monte Carlo (rjMCMC) method to collapse or split nodes in the guide species tree according to node posterior probabilities. We created bpp input files from the ‘.loci’ output file from pyrad using the R scripts *bpp_convert_Ama_sp.r* written by J-P. Huang and available at https://github.com/airbugs/Dynastes_delimitation (Huang 2018). We discarded loci that were not represented in at least one individual per taxon (i.e., loci with missing taxa were removed; e.g., Huang et al. 2020). The final dataset retained 333 loci. We considered as prior settings: *ϑ* = G (3, 0.002) and *τ* = G (3, 0.004), where *ϑ* and *τ* refer to the ancestral population sizes and divergence times, respectively. We ran two replicates and used an automatic adjustment of the finetune parameters, allowing swapping rates to range between 0.30 and 0.70 (Yang 2015). We ran each analysis for 100,000 generations, sampling every 2 generations (10,000 samples), after a burn-in of 50,000 generations. We evaluated convergence of replicates using tracer version 1.7.1 (Rambaut et al. 2018).

### Phylonetwork Reconstruction

Phylogenetic reconstruction without considering the potential occurrence of post-divergence gene flow (i.e., introgressive hybridization) can have severe impacts on the obtained inferences (Solís-Lemus and Ané 2016; Burbrink and Gehara 2018; Olave and Meyer 2020). Although the two phylogenomic inference methods employed (snapp and bpp) yielded the same most supported topology (see Results section), unsupported nodes led us to investigate the presence and impact of multiple branches connections using the julia package phylonetworks (Solís-Lemus et al. 2017). This method uses a maximum pseudolikelihood estimator applied to quartet concordance factors (CF) of 4-taxon trees under the coalescent model, incorporating incomplete lineage sorting and reticulation events (Solís-Lemus et al. 2017). The observed CF from the estimated gene trees is then used to estimate a semi-directed species network with estimated reticulation events and *γ*-values indicating the proportion of ancestral contribution to the hybrid lineage genome.

To estimate individual gene trees for each locus, we followed magnet version 0.1.5 pipeline (J. C. Bagley, http://github.com/justincbagley/MAGNET). We ran magnet pipeline using as input file the aligned DNA sequences from the pyrad output file ‘.gphocs’. Specifically, magnet first splits each locus contained in the ‘.gphocs’ file into separated phylip-formatted alignment files, and sets up and runs raxml (Stamatakis 2014) to infer a maximum-likelihood (ML) gene tree for each locus. Prior to obtain the gene trees, we applied trimal version 1.2 (Capella-Gutiérrez et al. 2009) to our phylip dataset in order to filter out loci with a high average identity (>0.99 %) across the multisequence alignment and retain only those that are most informative (Bernardes et al. 2007). Then, we used phylonetworks to read all raxml gene-trees retained (20,637 trees) and calculate CFs, with all individuals per clade mapped as alleles to species. We used the bpp/snapp tree as the starting topology, and tested values for *h* (number of reticulations) from 0 to 5, assessing maximum support using a slope heuristic for the increase in likelihood plotted against *h* (Solís-Lemus and Ané 2016). We ran 50 independent runs per *h*-value to ensure convergence on a global optimum.

### Divergence Time Estimation

We ran bpp under module A00 to obtain the posterior distribution of species divergence times (*τ*s) under the multispecies coalescent (MSC) model (Yang 2002; Rannala and Yang 2003). A recent implementation of A00 analysis on bpp version 4 allows estimating parameters under the MSC considering past introgression events (*φ*s) (multispecies coalescent with introgression, MSCi; Rannala and Yang 2003; Burgess and Yang 2008; Flouri et al. 2019). To evaluate the impact of introgression events on divergence time estimation, we conducted A00 analyses under both the MSC and MSCi models using as fixed topology i) the one most supported by snapp and A01 bpp analyses (MSC model) and ii) the species tree from the most supported phylogenetic network recovered using phylonetworks (MSCi model). For each analysis, we executed two runs and assigned values for the inverse-gamma priors *ϑ* ~ IG(3, 0.004) for all *ϑ* s and *τ* ~ IG(3, 0.004) for the age *τ*_0_ of the root as suggested in Flouri et al. (2019) when no information is available about prior parameters. A total of 50,000 iterations (sample interval of 5) with a burn-in of 10,000 was implemented for each run and convergence was evaluated across replicates using tracer (Rambaut et al. 2018). Divergence times were calculated according to the equation *t* = τ/2μ(e.g., Huang et al. 2020), where τ is the divergence in substitutions per site estimated by BPP, μ is the per site mutation rate per generation, and *t* is the absolute divergence time in years. We assumed a genomic mutation rate of 2.8 × 10^-9^ per site per generation (Keightley et al. 2014) and a one-year generation time (Clemente et al. 1990).

### Geometric Morphometric Analyses

To characterize phenotypic variation, we chose traits that have been used to delineate taxonomic units in the complex (pronotum; Clemente et al. 1991) and associated to courtship behavior (forewing; e.g., Nattier et al. 2011) and reproduction (male genitalia; e.g., Huang et al. 2020) in Orthoptera. We selected 10 individuals from each studied population (5 males and 5 females for each of the two populations per species/lineage, when available) to analyze forewing and pronotum variation and two individuals per population to extract and characterize male genitalia (penis lateral valve shape). To prepare male genitalia, we made a longitudinal cut and peeled back the apex of the abdomen to remove the exoskeleton. Abdominal contents were removed with fine forceps and placed in a Petri dish with 20% KOH for ~2 hours at room temperature to digest connective tissues. After that time, the sclerotized structure of the genitalia became apparent in the materials. We used landmark-based geometric morphometric methods (GMM) to characterize phenotypic variation in the selected traits. We captured digital images of dorsal views of pronota and forewings and of lateral views of male internal genitalia with a Leica MZ16 A stereomicroscope fitted with a DFC 450 camera using the Leica Application Software (LAS) version 3.8 (Leica Microsystems Ltd, Switzerland). We used fixed landmarks to characterize pronotum (9 landmarks) and forewing (11 landmarks) shape and a combination of fixed landmarks (3 landmarks) and semi-landmarks (35 landmarks) to capture the shape of male genitalia. Landmarks were mapped on the images using tpsdig version 2.2 (Rohlf 2015) and analyzed as implemented in the R version 3.3.2 (R Core Team, 2018) package geomorph (Adams and Otárola-Castillo 2013). Semi-landmarks were resampled to be equidistant along their curves and “slid” via minimizing bending energy (Bookstein 1992; Bookstein et al. 1999; Gunz et al. 2005). We obtained shape variations for each sex and trait through generalized Procrustes analyses (GPA) (Rohlf and Slice 1990; Rohlf 1999) with the package geomorph. Specifically, we performed GPA to standardize the size and remove the effects of location and rotation of the relative positions of landmarks among specimens using the function *gpagen*. This superimposition method minimizes the sum-of-squared distances between landmarks across samples (Rohlf and Slice 1990). We used principal components analysis (PCA) of the Procrustes coordinates for each dataset to extract the most explanatory axes of shape variation. To test for shape differences among species, we performed a Procrustes ANOVA using distributions generated from a resampling procedure based on 1,000 iterations in the R package geomorph using the function *procD.lm* (Adams and Collyer 2018). Significance values (*p*-values) between each pair of species were determined for each sex and trait using the *pairwise* function. To visualize shape differences, we represented the first two principal component axes (the most explicative) in a convex hull for each species and sex, using *ddplyr* function in the R package plyr (Wickham et al. 2019).

We quantified the phylogenetic signal (i.e., how morphologically similar closely related species are to one another) for each trait using Blomberg et al.’s (2003) *K* under the function *physignal* in geomorph. We used the tree topology most supported by the phylogenetic inference analyses detailed above and performed 1,000 permutations of shape data among the tips of the phylogeny to evaluate statistical significance. A *K*-value of 1 reflects perfect accord with expected patterns of shape variation under Brownian motion, values greater than 1 reflect phylogenetic under-dispersion of shape variation (i.e., close relatives are more similar than expected under Brownian motion), and values less than 1 indicate phylogenetic over-dispersion of shape variation (i.e., close relatives are less similar than expected under Brownian motion).

## Results

### Genomic Data

Illumina sequencing returned an average of 2.89 × 10^6^ reads per sample. After quality control, an average of 2.13 × 10^6^ reads was retained per sample (Supplementary Fig. S1 available on Dryad). The genomic datasets obtained with pyrad (*minCov* = 25%) for the subsets of 18 and 36 individuals retained a total of 50,192 and 21,438 variable loci, respectively.

### Phylogenomic Inference

Species trees reconstructed by snapp and bpp yielded the same topology and the two analyses only differed in the degree of support for some clades (Figs. 1 and 2). These analyses recovered three main monophyletic groups: a Maghrebian clade (*O. alluaudi* and *O. lepineyi*), an Iberian clade (*O. antigai* and *O. femoralis*), and an Ibero-Maghrebian clade (*O. lecerfi*, *O. bolivari*, *O. uhagonii* and *O. minutissimus*). The Maghrebian species *O. alluaudi* and *O. lepineyi* constituted the most basal and the Iberian and Ibero-Maghrebian clades shared a sister relationship (Figs. 1 and 2). snapp analyses showed low clade support (i.e., posterior probabilities values < 0.95) for internal relationships within the Ibero-Maghrebian clade. Accordingly, the most frequently recovered topology with snapp (37.84%) differed from the alternative less supported topologies on the sisterhood relationships among species within this group (Figs. 1 and 2). In contrast to snapp, posterior probabilities in bpp were consistently high for all clades, except for the split between *O. uhagonii* and the two *O. minutissimus* lineages (Fig. 2).

**Figure 2.**
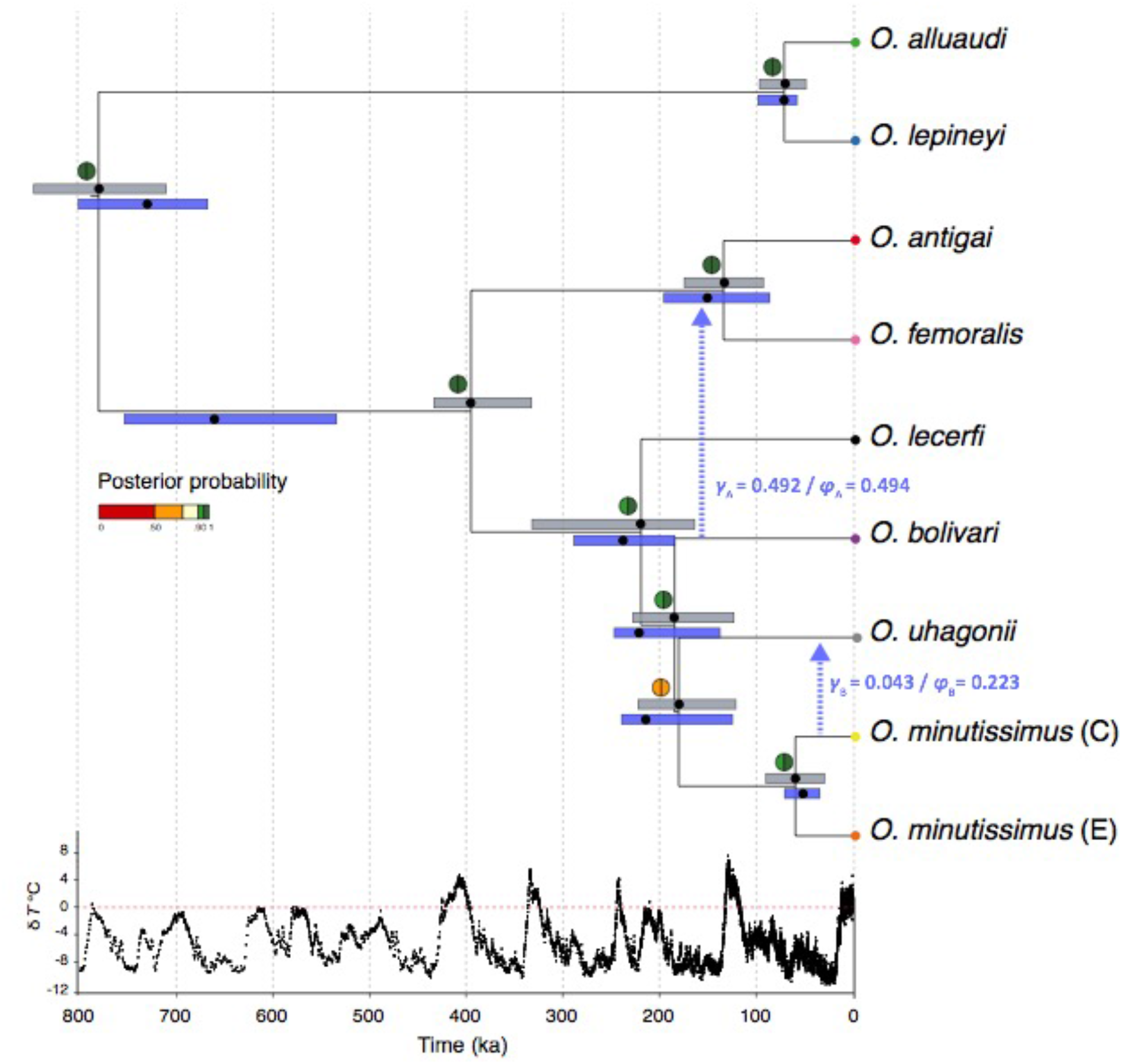
Species tree including estimates of divergence time and inferred introgression events. The species tree was reconstructed with snapp and bpp (option A01) and posterior probabilities of node support are indicated for each analysis in colored semi circles (left: snapp; right: bpp). Dots (median) and bars (95% highest posterior density intervals) indicate divergence times estimated by bpp (option A00), colored in grey for standard analysis not considering post-divergence gene flow (MSC model) and blue for analyses accounting for introgression events (MSCi model) inferred using phylonetworks. Blue arrows indicate inferred introgression events with their corresponding inheritance values (γ) estimated by phylonetworks and introgression probabilities (φ) estimated by bpp (not time-scaled). Bottom panel shows temperature anomaly (δ*T* °C) in the Late Quaternary as estimated from the EPICA (European Project for Ice Coring in Antarctica) Dome C ice core (Jouzel et al. 2007).

### Phylonetwork Reconstruction

phylonetwork analyses revealed that all models involving reticulation events (*h*> 0) fit our data better than models considering strict bifurcating trees (*h* = 0) (Supplementary Fig. S2 available on Dryad). The best phylogenetic network inferred by phylonetworks identified two introgression events (*hmax* = 2, negative pseudolikelihood = −6.20) and a backbone tree in concordance with the topologies recovered by snapp and bpp (Figs. 1 and 2). The optimal network supported introgression from *O. minutissimus*-C to the sympatric *O. uhagonii* (*γ*_A_ =0.043) and from *O. bolivari* to the most recent common ancestor (MRCA) of *O. femoralis* and *O. antigai* (*γ*_B_ =0.492) (Fig. 2).

### Divergence Time Estimation

Divergence times estimated by bpp both considering (MSCi model) and not considering (MSC model) post-divergence gene flow are summarized in Fig. 2. Both analyses supported that the initial split of the Maghrebian clade from the rest of the species took place during the Middle Pleistocene (~ 675 to 850 ka). Estimates of divergence time between the Iberian and the Ibero-Maghrebian clades was the most important discrepancy between the results yielded by bpp analyses under the MSC and MSCi models. bpp analyses not considering post-divergence gene flow estimated that these two clades split around 340-445 ka. However, divergence times obtained under the MSCi model yielded older estimates, around 540-770 ka. The 95% highest posterior density (HPD) intervals obtained under the two models largely overlapped for the rest of the nodes. Analyses showed that all contemporary species originated in the last ~200 ka, during the end of the Middle and the beginning of the Late Pleistocene (Fig. 2). Introgression from *O. minutissimus* C to *O. uhagonii* took place around 49 ka, whereas introgression from *O. bolivari* to the ancestor of *O. antigai* and *O. femoralis* dated back to 205 ka. The introgression probability (φ; Flouris et al. 2020) estimated by bpp for the introgression event from *O. bolivari* to the ancestor of *O. antigai* and *O. femoralis* was virtually identical (φ_A_ = 0.494) to the inheritance parameter (γ; Solís-Lemus and Ane 2016) estimated by phylonetwork (γ = 0.492; Fig. 2). However, the introgression probability from *O. minutissimus* C to *O. uhagonii* estimated by bpp was much higher (φ_A_ = 0.223) than the analogous inheritance parameter (γ = 0.043) yielded by phylonetwork analyses (Fig. 2).

### Analyses of Phenotypic Variation

A high proportion of pronotum and forewing shape variation was explained (>65%) by the first two principal components (Fig. 3). All analyzed traits significantly differed among species/lineages in both sexes (Supplementary Table S1 available on Dryad). In males, two extreme forms could be distinguished in forewing shape variation: a spindle-like shape (*O. lecerfi*) and an elongated trapezoid shape (*O. uhagonii*) (Fig. 3A). In females, species/lineages could be differentiated by rounded (*O. antigai* and *O. bolivari*), sharped (*O. alluaudi*, *O. lepineyi*, *O. femoralis* and *O. minutissimus* E) and intermediate (*O. uhagonii, O. lecerfi*, *O. minutissimus* C) forewing shapes (Fig. 3C). Forewing shape in the two sexes was significantly different in most pair-wise species/lineage comparisons (Supplementary Table S2 available on Dryad). Although dorsal pronotal shape variation in both males and females showed highly significant differences among species/lineages (Supplementary Table S1 available on Dryad), this trait tended to present a higher overlap than forewing shape variation (Fig. 3B, D). Accordingly, a fewer number of pair-wise species/lineage comparisons were statistically significant (Supplementary Table S3 available on Dryad). The lower number of samples analyzed for male genitalia made not possible a visual representation of shape variation for this trait. However, results from procrustes ANOVA showed significant differences among species/lineages that were mostly driven by differences between *O. lepineyi* and *O. alluaudi* (hooked shape) and the rest of the species (straight shape) (Supplementary Table S4 available on Dryad).

**Figure 3.**
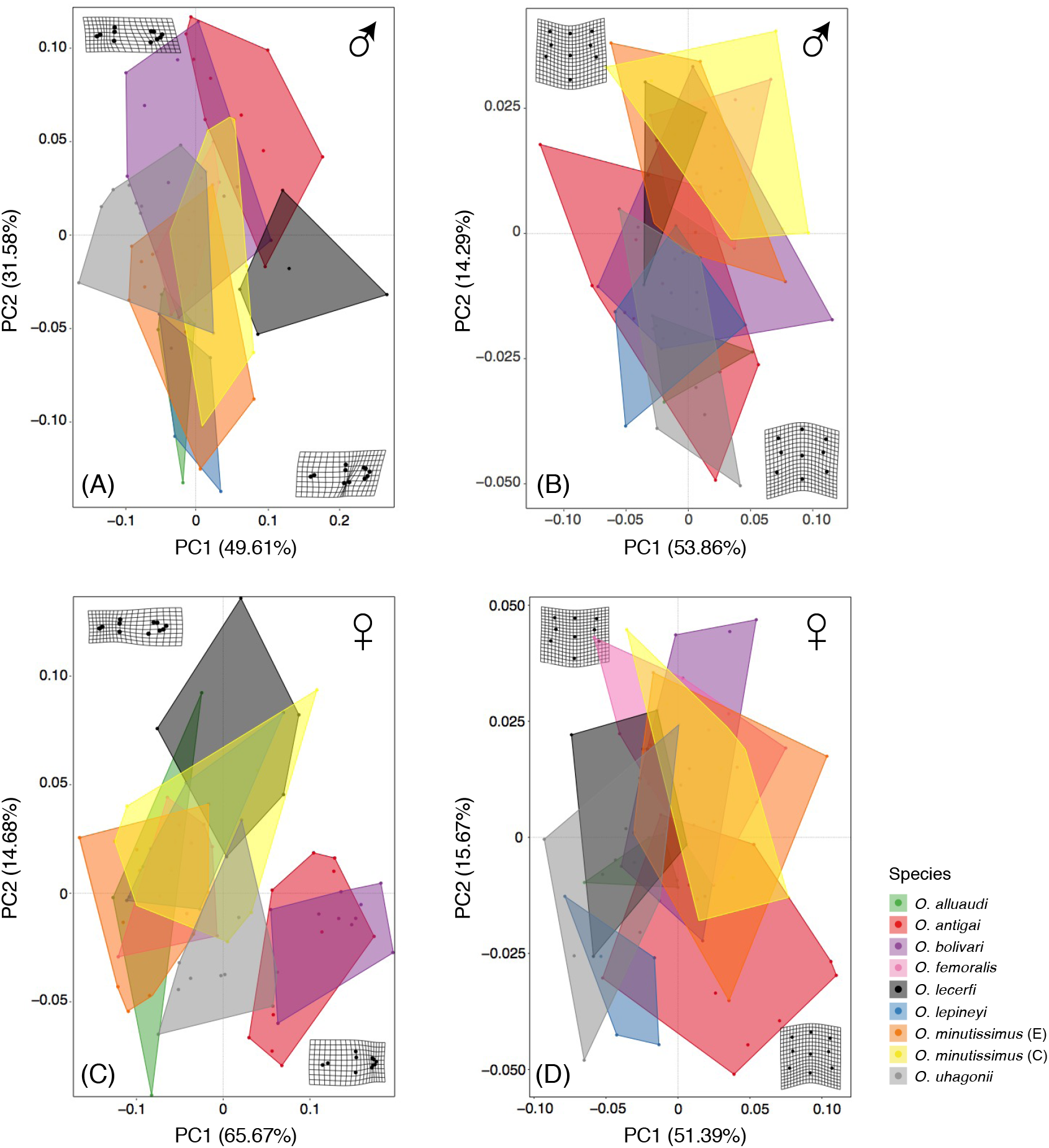
Principal component analyses for (A, C) forewing and (B, D) pronotum size-corrected shape variation in (A, B) males and (C, D) females. Colored convex hull polygons show species/lineage variation and warp grids represent extreme shape variation for the first two principal components.

In males, forewing, pronotum and genitalia shapes exhibited significant phylogenetic signals and *K* values < 1 indicated that closely related taxa are less similar in these traits than expected under Brownian motion (Fig. 4). The degree of phylogenetic signal varied across male traits, being weaker for male genitalia (Fig. 4). In females, forewing and pronotum shapes did not show a significant phylogenetic signal, albeit pronotum shape was marginally non-significant (Fig. 4).

**Figure 4.**
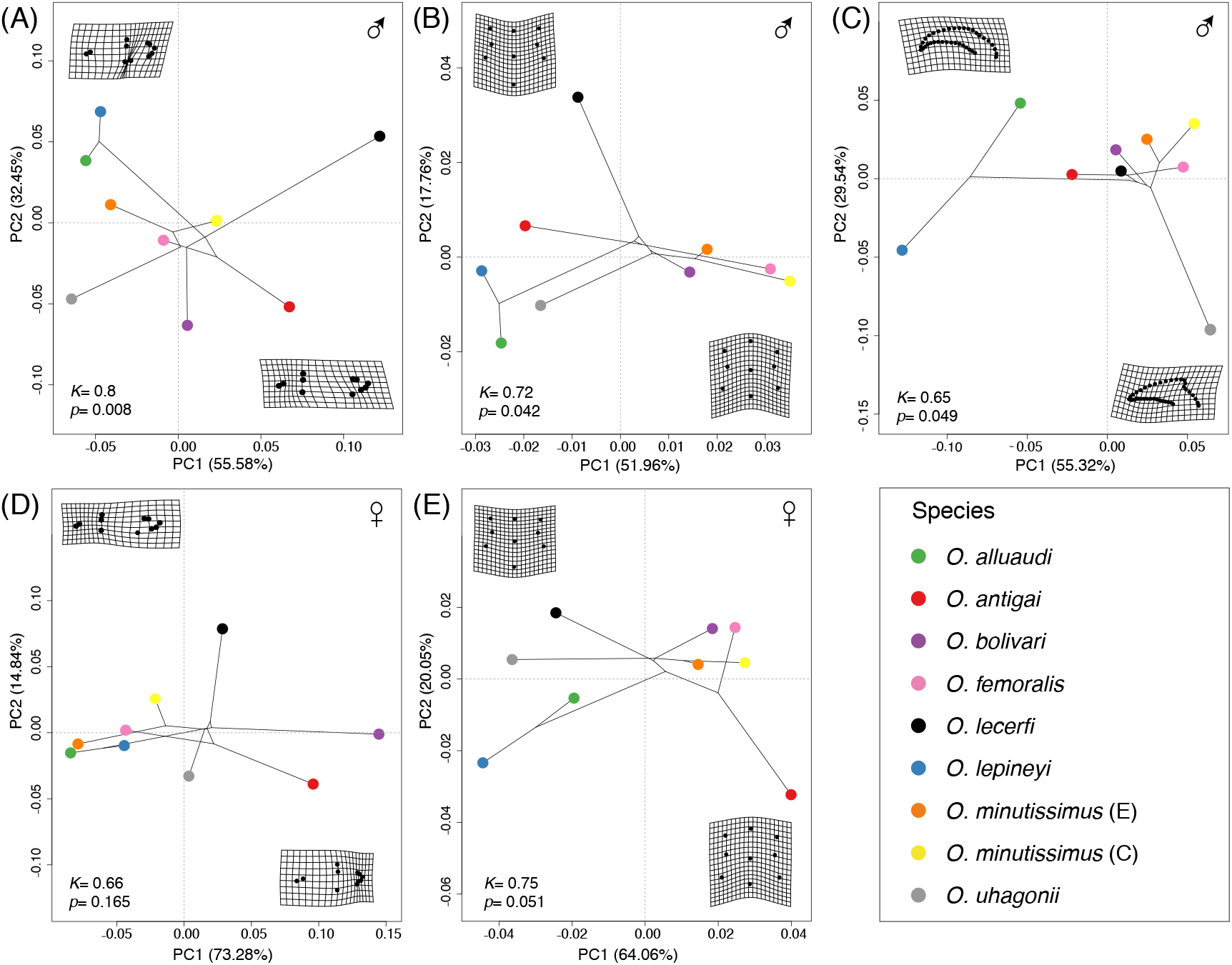
Phylomorphospaces showing the first two principal components (PC1 and PC2) from a PCA summarizing size-corrected shape variation for (A) forewing, (B) pronotum and (C) genitalia in males and (D) forewing and (E) pronotum in females. Colored dots indicate the different species and warp grids represent extreme shape variation for the first two principal components.

## Discussion

By reconstructing lineage and phenotypic diversification in a complex of montane grasshoppers, our study contributed to shed light on the ecological and evolutionary processes underlying the high rates of local endemism of the Ibero-Maghrebian biodiversity hotspot (Hewitt 1996; Avise and Wollenberg 1997). The combination of genomic data and a comprehensive suite of coalescent-based phylogenetic analyses provided strong support for a recent radiation (< 1 Ma) of the subgenus *Dreuxius*, indicating that periods of isolation and secondary contact linked to Pleistocene glacial cycles likely contributed to both allopatric speciation and post divergence gene flow. Geometric morphometric analyses for traits of taxonomic relevance and putatively involved in different components of reproductive isolation (sexual selection, copulation, etc.) supported the phenotypic distinctiveness of most sister taxa within the complex. Moreover, some of the studied traits presented a significantly lower phylogenetic signal than expected under a Brownian motion model of evolution, suggesting that phenotypic variation might have been in part shaped by natural or sexual selection acting at different stages of speciation (Kelly 2014; Servedio and Boughman 2017). This research exemplifies how the integration of multiple lines of evidence can help to reconstruct complex histories of reticulated evolution linked to Late Quaternary climatic changes and highlights the importance of implementing new methodological approaches to deal with post-divergence gene flow, a necessary step toward getting unbiased estimates of key demographic parameters and drawing a more realistic evolutionary portrait of Pleistocene radiations in which incomplete lineage sorting often co-occurs with introgressive hybridization (Wen et al. 2016; Nevado et al. 2018).

### Evolutionary Biogeography of the Species Complex

Our phylogenetic reconstructions and estimates of divergence time supported that the diversification of the subgenus *Dreuxius* took place over the last 800 ka, with direct ancestors of extant species tracing back their origins to end of the Middle Pleistocene (<250 Ka) (Fig. 2), and most splitting events occurring in a short time span (~200 ka). The earliest split within *Dreuxius* (ca. 800 ka) separated the lineage including the Maghrebian *O. alluaudi* and *O. lepineyi* from the most speciose clade including all Iberian taxa plus the northwestern African *O. lecerfi*. The last clade subsequently split (ca. 400 ka) into two clades, one formed by the Pyrenean *O. antigai* and the Baetican *O. femoralis* and another comprising the rest of Iberian species and *O. lecerfi*. Our genomic data support *O. antigai* and *O. femoralis* as sister taxa and a close relationship between *O. bolivari, O. minutissimus* and *O. uhagonii*, which agrees with previous descriptive assessments of species relationships based on morphological and behavioral comparisons (Gangwere and Morales Agacino 1970; Clemente et al. 1991). Our analyses also supported that the divergence between the two allopatric lineages of *O. minutissimus* is of the same order of magnitude (ca. 60 ka) than that estimated between the sister taxa *O. alluaudi* and *O. lepineyi* (Fig. 2). The genotypic and phenotypic distinctiveness of these two lineages (Cáliz 2015; Tonzo et al. 2020) call upon a taxonomic re-assessment of this monotypic taxon that was formerly composed by two distinct taxa: *O. burri* Uvarov, 1936 widely distributed in eastern Iberia and *O. minutissimus* (Brullé, 1832) restricted to the Central System (Clemente et al. 1990 and references therein).

The Pleistocene origin of all clades and lineages within the studied species complex points to the important role of Quaternary climatic oscillations in the transcontinental diversification of Ibero-Maghrebian biotas. With the exception of *O. minutissimus*, which is distributed from sea level to alpine areas above the tree line, the rest of taxa within the complex are montane species restricted to high elevations (>1,300 m) in different ranges from the region. Thus, Late Pleistocene climatic oscillations, when most speciation events within the complex took place, are expected to have contributed to create multiple opportunities for both divergence and post-divergence gene flow through elevational and latitudinal range-shifts (Hewitt 2000; Knowles 2000). Despite the genetic and phenotypic distinctiveness of the species within the complex and their current distribution in distant mountain ranges, the habitats occupied show strong similarities across all taxa (Ragge 1986; Clemente et al. 1990; Clemente et al. 1991; Tonzo et al. 2020). This points to allopatric speciation, rather than ecological divergence, as the predominant mechanism of species diversification (Taberlet et al. 1998; Hewitt 2000; Hewitt 2004; Mayer et al. 2010). Topographically complex regions such as Iberia and northwestern Africa offer an ideal biogeographic setting for allopatric speciation, as isolation in valleys during glacial periods (i.e., glacial refugia; Knowles 2001; Wallis et al. 2016) and confinement in sky-islands during interglacials (i.e., interglacial refugia; Bennett and Provan 2008; Stewart et al. 2010) are expected to lead to extended periods of isolation and divergence through genetic drift and/or natural selection under contrasting selective regimes (Hewitt 1996; Djamali et al. 2012). Furthermore, range shifts might have contributed in some cases to complete the speciation process through the evolution of reproductive isolation in secondary contact zones (i.e., reinforcement; Butlin 1989, 1998; Hewitt 2008; Tonzo et al. 2020).

Phylogenetic analyses did not recover Maghrebian taxa as a monophyletic clade, supporting two trans-continental colonization events through the Strait of Gibraltar or adjacent areas. Glacial periods reduced the Mediterranean Sea level about 125 m and shortened the distance between northwestern Africa and southern Iberia to less than 5 km, which might have led to the emergence of small islands and shoals and facilitated the exchange of biotas between the two continents during the coldest stages of the Pleistocene (Collina-Girard 2001; Cosson et al. 2005; Agustí et al. 2006). These results add to the accumulating empirical evidence supporting the migration of numerous organisms across the two continents, either seeking for glacial refugia in North Africa or following post glacial colonization routes to Europe (Taberlet et al. 1998; Teacher et al. 2009; Graciá et al. 2013; Husemann et al. 2014).

### A Reticulated Evolutionary History

Interspecific gene flow and ILS are ubiquitous phenomena in recent evolutionary radiations and, thus, require to be evaluated when inferring phylogenetic relationships and demographic parameters in species complexes of Pleistocene origin (Yu and Nakhleh 2015; Solís-Lemus and Ané 2016; Wen et al. 2018). Although the monophyly of the three main clades of the subgenus *Dreuxius* was consistently well-supported, internal nodes of the most speciose clade showed weak support in both bpp and snapp analyses (Fig. 2). Phylogenetic network analyses point to interspecific gene flow, rather than ILS, as the main cause of gene tree conflict (Fig. 2). Specifically, we found two events of introgression involving lineages at different stages of the speciation continuum: from *O. bolivari* to the most recent common ancestor of *O*. *antigai* and *O*. *femoralis* (ca. 205 ka) and from the lineage of *O. minutissimus* distributed in the Central System to its sympatric counterpart *O*. *uhagonii* (ca. 49 ka) (Fig. 2). As expected, the two introgression events involved taxa from the same continental landmass (i.e., Iberian Peninsula). *Omocestus bolivari* and *O. femoralis* currently present adjacent but non-overlapping distributions in the sky island archipelago of the Baetic System (Fig. 1). However, genomic-based demographic inferences have recently revealed that the two species experienced considerable expansions during the last glacial period, when their ranges likely overlapped according palaeodistribution reconstructions (V. Tonzo and J. Ortego, in prep.). This is expected to have led to secondary contact and might explain the detected signatures of historical gene flow from *O. bolivari* to the common ancestor of *O. femoralis* and *O. antigai*. The very low support for the split between *O. minutissimus* and *O. uhagonii* was explained by historical hybridization between the two taxa in the Central System, where the evolution of reproductive isolation via reinforcement or other mechanisms has been hypothesized to prevent gene flow among contemporary sympatric populations of the two species (Tonzo et al. 2020).

Although there is an increasing interest on implementing phylogenomic network approaches to empirical data (Eckert and Carstens 2008; Pickrell and Pritchard 2012; Yu and Nakhleh 2015; Solís-Lemus et al. 2017; Wen et al. 2018), the impact of interspecific gene flow on inferred divergence times has been rarely evaluated (Flouri et al. 2019). We assessed the impact of introgression on the estimated timing of species split and found that, as expected, ignoring interspecific gene flow result in an underestimation of divergence times in some nodes. Specifically, the timing of divergence between *O. antigai*-*O. femoralis* and the rest of the species was estimated to be ca. 200 ka older when analyses accounted for introgression, whereas historical gene flow between sympatric populations of the more recently diverged *O. minutissimus* and *O. uhagonii* had a little impact on our inferences.

### Phenotypic Variation

We found that all the studied phenotypic traits differed among lineages, with most species/lineage pairs presenting significant differences in at least one of them (Supplementary Tables S1-4 available on Dryad). In both sexes, forewing shape tended to show stronger differences among species than pronotum and male genitalia (Fig. 3 and Supplementary Tables S1-4 available on Dryad). Forewings are involved in courtship acoustic behavior in grasshoppers (Von Helversen et al. 2004; Vedenina and Mugue 2011; Ronacher 2019), a character directly implicated in mate attraction and subjected to sexual selection (Oh and Shaw 2013; Outomuro et al. 2016). Traits under sexual selection can evolve rapidly, accelerating speciation when other forces as ecological adaptations are not so evident or absent (Anderson 1994; Mendelson and Shaw 2005; Rundell and Price 2009). Accordingly, species within the *Dreuxius* species complex show very similar habitat requirements but present distinctive songs (Ragge 1986; Reynolds 1987; Clemente et al. 1991, 1999), which suggests that sexual selection might have played an important role in the completion of the speciation process (e.g., Bridle and Butlin 2002; Bridle et al. 2006) and prevented interbreeding among contemporary sympatric populations in secondary contact zones (Tonzo et al. 2020). In the case of male genitalia, we found that interspecific variation was mostly determined by differences between the two earliest diverged clades (Supplementary Table S4 available on Dryad). Although differences among species in this trait must be interpreted with extreme caution due to small sample sizes, the fact that some currently sympatric lineages (e.g., *O. minutissimus* and the rest of Iberian species) share similar male genitalia suggests that reproductive isolation might have been driven by other phenotypic or behavioral traits (e.g., mate selection). Remarkably, species/lineages involved in historical introgression presented significant phenotypic differences for one or more of the studied traits, suggesting that historical hybridization has not led to phenotypic assimilation (Huang 2016) or that, on the contrary, secondary contact might have contributed to phenotypic divergence through some form of character displacement (Pfennig and Pfennig 2009). Finally, phylomorphospace analyses and the *K* statistic of Blomberg et al. (2003) (*K*<1 in all cases) indicated that species are less similar at some of the studied morphological traits than expected under a Brownian motion model of evolution (Fig. 4). Even when phylogenetic signal alone is not a direct way of elucidating the evolutionary processes responsible for phenotypic diversification, these results also suggests that natural and/or sexual selection might have modulated phenotypic diversification in the complex (Blomberg et al. 2003; Pennell and Harmon 2013).

## Conclusions

Our study exemplifies the importance of integrating different sources of information to reconstruct complex biogeographic histories and understand the processes underlying the high rates of local endemism in the Ibero-Maghrebian transcontinental biodiversity hotspot. Although the retrieved topology, estimates of divergence time (i.e., Pleistocene) and biogeographic inferences did not qualitative change considering or not inter-specific gene flow, past hybridization events are an important component of speciation that must be resolved to shed light on the evolutionary pathways of recent species complexes. Collectively, our analyses demonstrate a very recent origin of the studied radiation (< 1 Ma) and support the permeability of the Strait of Gibraltar to the exchange of low-vagile terrestrial fauna during the Pleistocene (Husemann et al. 2014), rejecting the hypothesis of a protracted history of divergence dating back to ancient southern Europe-northern Africa connections during the Tortonian or the Messinian (e.g., Hidalgo-Galiana and Ribera 2011; Faille et al. 2014; Ortego et al. 2017). This points to the important impact of Pleistocene glaciations as a diversification engine in the Ibero-Maghrebian region, which has been often assumed to have been scarcely impacted by Quaternary glaciations due to its low latitude and temperature buffering by the Atlantic Ocean and the Mediterranean Sea (Rodríguez-Sánchez et al. 2008).

## Supplementary Material

Data available from the Dryad Digital Repository: https://datadryad.org/stash/share/TyAdIXuDe8IBEWGmZ7ULxR3a14OSCPGEWRgOnzwPSgA. Raw Illumina reads have been deposited at the NCBI Sequence Read Archive (SRA) under BioProject PRJNA543714.

## Acknowledgements

We are much indebted to Anna Papadopoulou for her valuable help in study design and useful comments, suggestions and corrections on a first draft of the manuscript. We are also grateful to Amparo Hidalgo-Galiana, Víctor Noguerales, and Pedro J. Cordero for their valuable help during field and laboratory work, Rosa Fernandez for her suggestions and help using trimal and Sergio Pereira (The Centre for Applied Genomics) for Illumina sequencing. Logistical support was provided by Laboratorio de Ecología Molecular (LEM-EBD) from Estación Biológica de Doñana. We thank to Centro de Supercomputación de Galicia (CESGA) and Doñana’s Singular Scientific-Technical Infrastructure (ICTS-RBD) for access to computer resources.

