## Supporting information for "Reticulate Evolutionary History in a Recent Radiation of Montane Grasshoppers Revealed by Genomic Data"

Data available from the Dryad Digital Repository:XXXXX

**Contents:**

Supplemental Methods

Methods S1. Genomic library preparation

Methods S2. Processing of genomic data

Supplemental Figures

Figure S1. Number of reads per individual before and after different quality filtering steps by pyrad and stacks

Figure S2. Pseudolikelihood score as a function of the number of reticulation events

Supplemental Tables

Table S1. ANOVA analyses testing differences among species/lineages in shape variation for the different studied traits

Table S2. Pairwise species/lineage comparisons for male and female forewing shape

Table S3. Pairwise species/lineage comparisons for male and female pronotum shape

Table S4. Pairwise species/lineage comparisons for male genitalia shape

Supplemental References

Supplemental Methods

Methods S1. *Genomic Library Preparation*

We used Nucleo Spin Tissue kits (Macherey-Nagel, Düren, Germany) to extract and purify total genomic DNA from a hind leg of each individual. Genomic DNA was individually barcoded and processed in house using the double-digest restriction-site associated DNA procedure (ddRADseq) described in Peterson et al., (2012), with minor modifications as detailed in Lanier et al. (2015) and Massatti and Knowles (2016). Briefly, DNA was doubly digested with EcoRI and MseI restriction enzymes, followed by the ligation of Illumina adaptor sequences and unique 7-base-pair barcodes. Ligation products were pooled into a library, size selected for fragments between 475 and 580 bp using a Pippin prep machine (Sage Science, Beverly, MA, USA), and amplified by iProofTM High-Fidelity DNA Polymerase (BIO-RAD, Veenendaal, The Netherlands) with 10-12 cycles. Single-read 151-bp sequencing was performed on an Illumina HiSeq2500 platform at The Centre for Applied Genomics (Hospital for Sick Children, Toronto, ON, Canada).

Methods S2. *Processing of Genomic Data*

Raw sequences were demultiplexed using *process_radtags*, a program distributed as part of the stacks pipeline (Catchen et al. 2011, 2013). Only reads with Phred scores ≥ 10 (using a sliding window of 15%), no adaptor contamination, and unambiguous barcode and restriction cut sites were retained. Read quality was checked in fastqc version 0.11.5 (A. Simon, <http://www.bioinformatics.babraham.ac.uk/projects/fastqc/>) and sequences were trimmed to 130 bp using seqtk (L. Heng, <https://github.com/lh3/seqtk>) to remove low-quality reads near the 3’ ends. The filtered and trimmed reads were assembled into de novo loci using pyrad version 3.0.66 (Eaton 2014). An additional quality-filtering step was performed with pyrad to convert base calls with a Phred score < 20 into Ns and discard reads with > 2 Ns. Parameter values for clustering threshold of sequence similarity (*W*_CLUST_ = 0.85), minimum coverage depth (*d* = 5), maximum individuals with shared heterozygous sites (*maxSH* = p.10), and maximum number of polymorphic sites in a final locus (*maxSNPs* = 20) were selected based on suggestions from the literature (Eaton and Ree 2013; Eaton 2014; Takahashi et al. 2014). Final datasets for subsequent analyses were generated discarding loci that were not present in at least ~25 % of the samples.

Supplemental Figures

Figure S1. Number of reads per individual before and after different quality filtering steps by pyrad and stacks. The cumulative stacked bars represent the total number of raw reads for each individual. Dark yellow color represents the reads that were discarded by *process_radtags* in stacks due to low quality, adapter contamination or ambiguous barcode. Red color represents the reads that were discarded during step 2 in pyrad after filtering out reads that did not comply with the quality criteria (reads with >2 sites with a Phred quality score < 20 were discarded). Green color represents the total number of retained reads used to identify homologous loci.

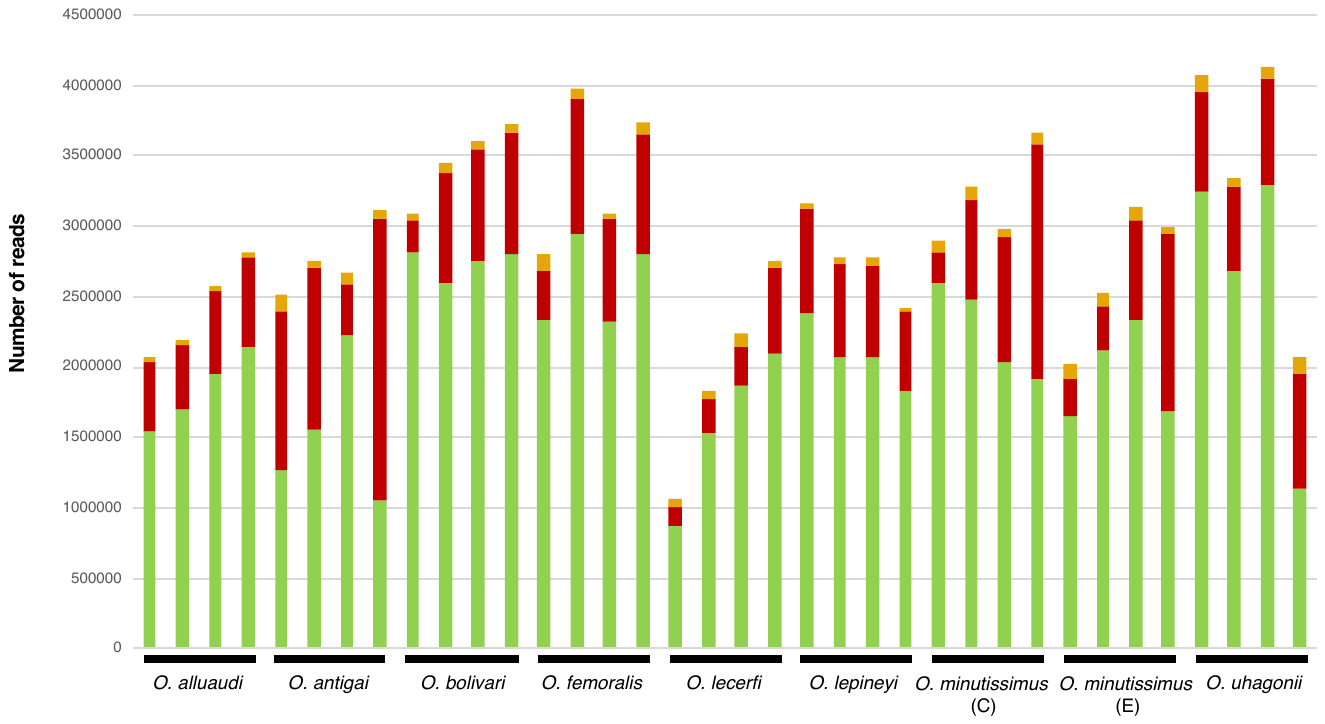

Figure S2. Plot of pseudolikelihood score as a function of the number of reticulation events to estimate network complexity in phylonetworks. Here, the slope heuristic suggests the presence of two hybrid nodes (*hmax* = 2).

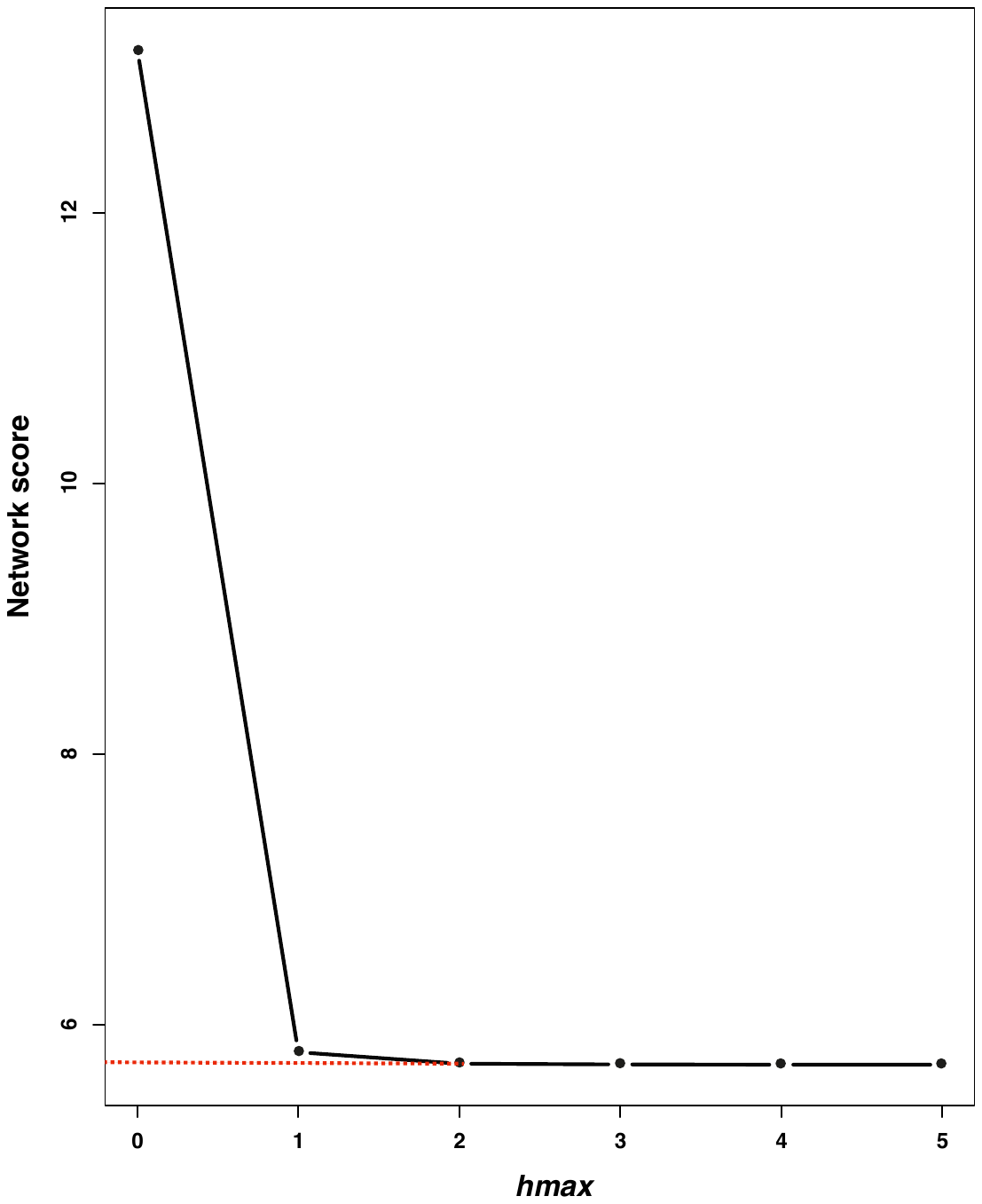

Supplemental Tables

Table S1. Results of ANOVA analyses testing differences among species/lineages in shape variation for the different studied traits. *Z*, effect sizes based on a *F* distribution; *P*, significance level.

| **Trait** | ***Z*** | ***P*** |
| --- | --- | --- |
| Male forewing | 6.724 | 0.001 |
| Female forewing | 7.177 | 0.001 |
| Male pronotum | 3.631 | 0.001 |
| Female pronotum | 5.226 | 0.001 |
| Male genitalia | 1.969 | 0.031 |

Table S2. Pairwise species/lineage comparisons between means of Procrustes shape for male (above the diagonal) and female (below the diagonal) forewing. Table shows *P*-values.

|  | *O. alluaudi* | *O. antigai* | *O. bolivari* | *O. femoralis* | *O. lecerfi* | *O. lepineyi* | *O. minutissimus* (E) | *O. minutissimus* (C) | *O. uhagonii* |
| --- | --- | --- | --- | --- | --- | --- | --- | --- | --- |
| *O. alluaudi* | - | 0.002 | 0.004 | 0.187 | 0.003 | 0.766 | 0.501 | 0.032 | 0.062 |
| *O. antigai* | 0.001 | - | 0.007 | 0.015 | 0.002 | 0.001 | 0.001 | 0.004 | 0.001 |
| *O. bolivari* | 0.001 | 0.098 | - | 0.060 | 0.001 | 0.002 | 0.012 | 0.050 | 0.034 |
| *O. femoralis* | 0.508 | 0.001 | 0.001 | - | 0.001 | 0.058 | 0.237 | 0.182 | 0.064 |
| *O. lecerfi* | 0.012 | 0.012 | 0.004 | 0.042 | - | 0.002 | 0.001 | 0.007 | 0.001 |
| *O. lepineyi* | 0.423 | 0.003 | 0.001 | 0.409 | 0.039 | - | 0.172 | 0.010 | 0.01 |
| *O. minutissimus* (E) | 0.767 | 0.001 | 0.001 | 0.389 | 0.005 | 0.110 | - | 0.075 | 0.078 |
| *O. minutissimus* (C) | 0.131 | 0.001 | 0.001 | 0.468 | 0.101 | 0.121 | 0.088 | - | 0.004 |
| *O. uhagonii* | 0.058 | 0.008 | 0.001 | 0.086 | 0.021 | 0.088 | 0.034 | 0.056 | - |

|  | *O. alluaudi* | *O. antigai* | *O. bolivari* | *O. femoralis* | *O. lecerfi* | *O. lepineyi* | *O. minutissimus* (E) | *O. minutissimus* (C) | *O. uhagonii* |
| --- | --- | --- | --- | --- | --- | --- | --- | --- | --- |
| *O. alluaudi* | - | 0.111 | 0.052 | 0.024 | 0.087 | 0.747 | 0.058 | 0.011 | 0.273 |
| *O. antigai* | 0.003 | - | 0.093 | 0.007 | 0.112 | 0.15 | 0.042 | 0.001 | 0.296 |
| *O. bolivari* | 0.071 | 0.012 | - | 0.252 | 0.069 | 0.054 | 0.388 | 0.152 | 0.111 |
| *O. femoralis* | 0.056 | 0.006 | 0.660 | - | 0.042 | 0.015 | 0.813 | 0.377 | 0.014 |
| *O. lecerfi* | 0.463 | 0.003 | 0.140 | 0.055 | - | 0.243 | 0.120 | 0.022 | 0.053 |
| *O. lepineyi* | 0.422 | 0.001 | 0.003 | 0.001 | 0.144 | - | 0.066 | 0.017 | 0.249 |
| *O. minutissimus* (E) | 0.163 | 0.019 | 0.648 | 0.655 | 0.118 | 0.009 | - | 0.231 | 0.043 |
| *O. minutissimus* (C) | 0.055 | 0.04 | 0.414 | 0.837 | 0.042 | 0.001 | 0.631 | - | 0.005 |
| *O. uhagonii* | 0.102 | 0.001 | 0.006 | 0.001 | 0.583 | 0.146 | 0.004 | 0.002 | - |

Table S3. Pairwise species/lineage comparisons between means of Procrustes shape for male (above the diagonal) and female (below the diagonal) pronotum. Table shows *P*-values.

|  | *O. alluaudi* | *O. antigai* | *O. bolivari* | *O. femoralis* | *O. lecerfi* | *O. lepineyi* | *O. minutissimus* (E) | *O. minutissimus* (C) | *O. uhagonii* |
| --- | --- | --- | --- | --- | --- | --- | --- | --- | --- |
| *O. alluaudi* | - | 0.678 | 0.756 | 0.170 | 0.575 | 0.147 | 0.366 | 0.220 | 0.010 |
| *O. antigai* |  | - | 0.931 | 0.475 | 0.960 | 0.146 | 0.743 | 0.446 | 0.104 |
| *O. bolivari* |  |  | - | 0.792 | 0.998 | 0.059 | 0.988 | 0.898 | 0.132 |
| *O. femoralis* |  |  |  | - | 0.878 | 0.004 | 0.649 | 0.937 | 0.210 |
| *O. lecerfi* |  |  |  |  | - | 0.061 | 0.976 | 0.831 | 0.227 |
| *O. lepineyi* |  |  |  |  |  | - | 0.012 | 0.003 | 0.003 |
| *O. minutissimus* (E) |  |  |  |  |  |  | - | 0.808 | 0.105 |
| *O. minutissimus* (C) |  |  |  |  |  |  |  | - | 0.107 |
| *O. uhagonii* |  |  |  |  |  |  |  |  | - |

Table S4. Pairwise species/lineage comparisons between means of Procrustes shape for male genitalia. Table shows *P*-values.
